# A fast-killing tyrosine amide ((*S*)-SW228703) with blood and liver-stage antimalarial activity associated with the Cyclic Amine Resistance Locus (*Pf*CARL)

**DOI:** 10.1101/2022.10.16.512381

**Authors:** Leah S. Imlay, Aloysus K. Lawong, Suraksha Gahalawat, Ashwani Kumar, Chao Xing, Nimisha Mittal, Sergio Wittlin, Alisje Churchyard, Hanspeter Niederstrasser, Benigno Crespo-Fernandez, Bruce Posner, Francisco Javier Gamo, Jake Baum, Elizabeth A. Winzeler, Benoît Laleu, Joseph M. Ready, Margaret A. Phillips

## Abstract

Current malaria treatments are threatened by drug resistance and new drugs are urgently needed. In a phenotypic screen for new antimalarials, we identified (*S*)-SW228703 ((*S*)-SW703), a tyrosine amide with asexual blood and liver stage activity and a fast-killing profile. Resistance to (*S*)-SW703 is associated with mutations in *Plasmodium falciparum* cyclic amine resistance locus (*Pf*CARL) and *P. falciparum* acetyl CoA transporter (*Pf*ACT), similarly to several other compounds that share features such as fast activity and liver-stage activity. Compounds with these resistance mechanisms are thought to act in the ER, though their target(s) are unknown. The tyramine of (*S*)-SW703 is shared with some reported *Pf*CARL-associated compounds; however, we observed that strict S-stereochemistry was required for activity of (*S*)-SW703, suggesting differences in mechanism of action or binding mode. (*S*)-SW703 provides a new chemical series with broad activity on multiple life-cycle stages and a fast-killing mechanism of action, available for lead optimization to generate new treatments for malaria.

---

The World Health Organization estimates that malaria caused 241 million infections and 627,000 deaths in 2020, with most deaths occurring among children in Africa.^1^ Widespread use of artemisinin combination therapies, as well as bed nets and other control measures, are credited with a decrease in case numbers and mortality since 2000, but the decline in malaria deaths has slowed in recent years. Further progress is threatened by growing resistance to artemisinins and other antimalarial therapies.^2^ Resistance to artemisinin (ART) and its analogs (e.g., artemether, artesunate) emerged and spread first in Southeast Asia;^3^ however, recent studies have also identified ART-resistant parasites in Africa, endangering malaria treatment and control programs in the most vulnerable region of the world.^4^ New treatments are urgently needed to combat the continuing threat of this devastating disease.

Malaria is transmitted by the bite of an infected mosquito, which injects sporozoite-stage parasites into the human host. These sporozoites migrate to the liver, invade hepatocytes, and differentiate into liver-stage schizonts.^5, 6^ Upon rupture, parasites invade erythrocytes (red blood cells; RBCs) and divide asexually, eventually rupturing the host cell and re-invading new RBCs. Malaria symptoms are caused by these intraerythrocytic stages, and thus drugs used to treat symptomatic malaria require intraerythrocytic activity. A minority of blood-stage parasites develop into gametocytes, which reproduce sexually upon uptake via mosquito bite, enabling the spread of the infection to a new host; compounds that are active against sexual stages block transmission and drugs that act against the liver stage are of interest for prophylaxis. ^7, 8^ Because of the small number of parasites during hepatic stages, it has been suggested that liver-active compounds may be less prone to resistance development if used for prophylaxis.^9^ Fast-acting compounds such as artesunate are also being prioritized for development, especially in cases of severe malaria, because they provide rapid relief of symptoms.^2^

Most antimalarials in current development were identified in phenotypic high-throughput screens (HTS) against the asexual RBC stages,^10, 11^ though more recent screens to identify compounds with liver stage^12^ and transmission blocking activity^13^ have also been reported. Target-based approaches have also been successfully used to identify clinical candidates. For example, in the case of DSM265, a dihydroorotate dehydrogenase (DHODH) enzyme-based high-throughput screen identified a hit compound, which was pursued in a structure-based lead optimization program, eventually identifying DSM265 as a clinical candidate.^6^ In contrast to target-based screens, phenotypic screening ensures that hit compounds will be cell-permeable,^11^ but subsequent target identification—helpful for lead optimization and toxicological profiling—can be challenging. In malaria drug discovery programs, resistance screening followed by whole genome sequencing has proven a robust mechanism to identify the targets of many compounds.^14^ For example, this approach was successfully used to identify the targets of several antimalarial candidates currently in development, including the clinical candidate KAE609 (NITD609/cipargamin),^15^ which targets ATP4; KDU691 and MMV390048, which target PI4K;^16, 17^ and several other phenotypic hits, which target tRNA synthetases ^18^ and acetyl CoA synthetase.^19^ Additionally, while the target remains unknown, similar studies have identified the *P. falciparum* cyclic amine resistance locus (*Pf*CARL), acetyl CoA transporter (*Pf*ACT), and UDP-galactose transporter (*Pf*UGT) as resistance proteins for imidazolopiperazines, including KAF156/ganaplacide, currently in stage IIb of clinical development.^20, 21^

Herein we describe SW228703 ((*S*)-SW703), a novel tyrosine amide scaffold with fast-killing activity against the blood stages of *P. falciparum*. (*S*)-SW703 was discovered as part of a phenotypic high-throughput screen (HTS) for compounds with asexual blood-stage antimalarial activity against *P. falciparum*, but was also found to have liver-stage activity. We selected for resistance to (*S*)-SW703 in *P. falciparum.* Subsequent whole genome sequencing (WGS) revealed that resistance is associated with mutations in *Pf*CARL and *Pf*ACT. (*S*)-SW703 thus represents a new chemical series that develops resistance via the multi-drug resistance locus, *Pf*CARL, further supporting the permissiveness of this resistance mechanism towards multiple unrelated scaffolds. (*S*)-SW703 has drug-like properties, is active only as the S-enantiomer, suggesting a protein target, and would serve as a strong starting point for medicinal chemistry if additional compounds with liver-stage activity are needed for antimalarial therapy.

## Results

### Identification of Hit Compound SW228703

A compound library of 100,000 small molecules was screened at 2.5 μM against asexual blood-stage *Plasmodium falciparum,* clone 3D7 (*Pf*3D7), using a previously published SyBR Green assay ^22^ optimized for our HTS core.^23^ The screen output was robust, with an average Z’ of 0.64. We found 1849 hits (RZ < −3), and the top 300 hit compounds (RZ score < −7.6) were retested at 3 concentrations (3, 1, and 0.3 μM) against both *Pf*3D7 and the human leukemia cell line, K562. Confirmed hits without cytotoxicity to K562 cells (CC_50_ > 3 μM) were ranked by potency versus *Pf*3D7 and filtered to remove previously discovered antimalarial compounds. Compounds of interest were further confirmed in more extensive dose-response experiments against *P. falciparum* and human HepG2 cells using newly-acquired compound (Chembridge Corporation and in-house synthesis). One of these compounds, SW228703 ((*S*)-SW703) (Fig. 1A), is a tyrosine amide that was found to have low-micromolar activity against *P. falciparum* asexual blood stages (3D7 EC_50_ = 0.78 μM; Dd2 EC_50_ = 0.81 μM) (Fig. 1B and Table 1) while showing no cytotoxicity versus a human cell line (HepG2 CC_50_ > 50 μM) (Table 1). (*S*)-SW703 has similar activity against a *P. falciparum* strain expressing DHODH from *Saccharomyces cerevisiae* (yDHODH), which bypasses the need for the mitochondrial DHODH and bc1 complex enzymes,^24^ ruling out both as targets of the compound. (*S*)-SW703 has good drug-like properties, conforming to Lipinski’s rule of 5,^25^ and it is highly soluble (> 34 μg/mL) at both pH 2 and 6.5 (Table 1). (*S*)-SW703 has poor metabolic stability in human liver microsomes (CL_int_ = 197 μL/min/mg protein), and, as with most early hit compounds, will require further optimization if the series is pursued.

**Figure 1.**
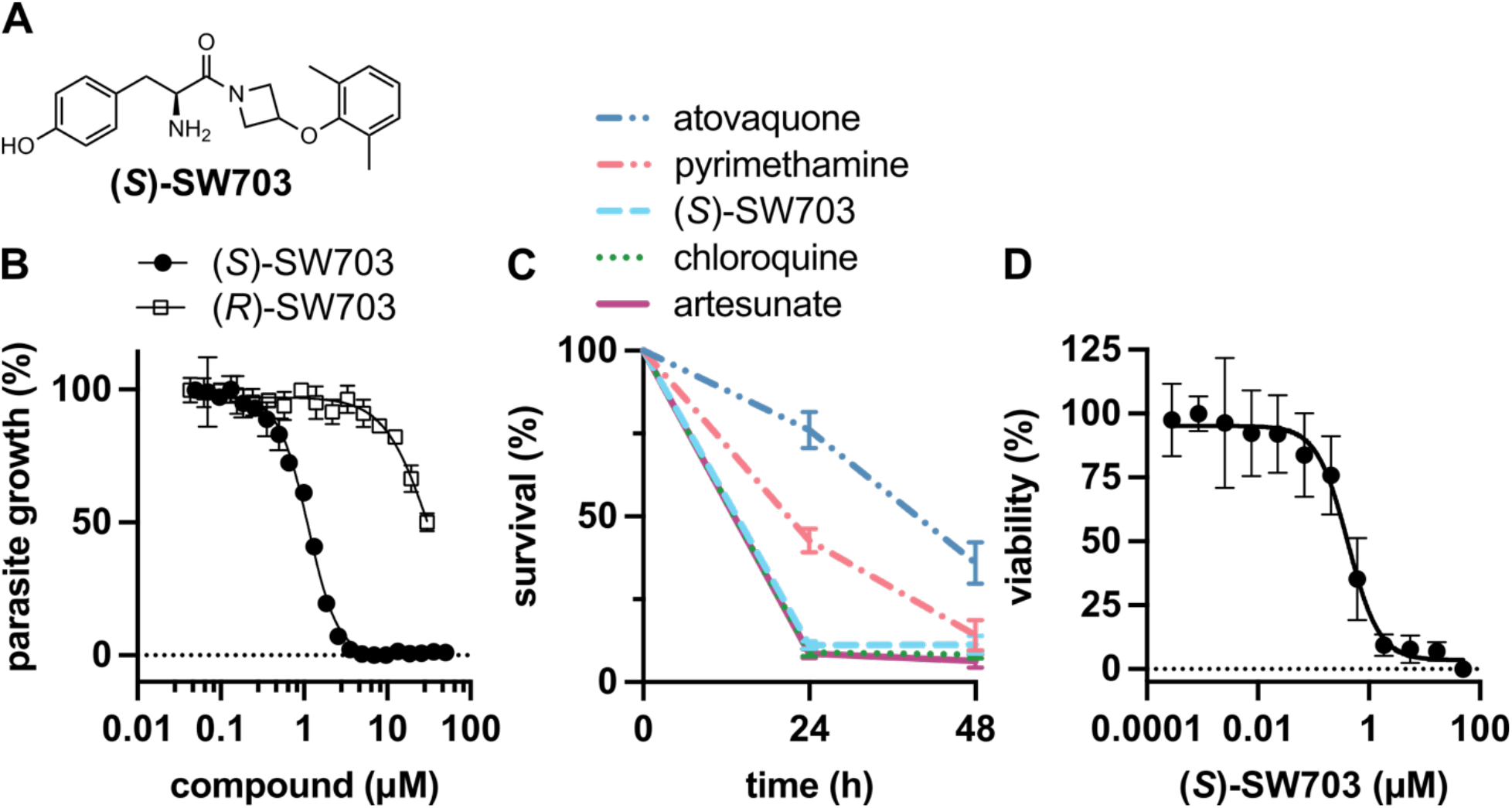
(*S*)-SW703 is a potent, enantiomerically specific, and fast-killing inhibitor of *P. falciparum* asexual blood-stages. (A) Structure of (*S*)-SW703. (B) *P. falciparum* (strain Dd2) growth inhibition (asexual intraerythrocytic parasites) by (*S*)-SW703 and its R-enantiomer; representative data from one study (3 technical replicates) shown; mean ± standard deviation. Table 1 shows the average data from at least three independent studies per compound. (C) To assess rate of kill, parasites were incubated with (*S*)-SW703 or benchmark compounds for 0, 24, or 48 hours at 10X EC_50_. Compounds were washed off and treated parasites were assessed for re-invasion of labeled erythrocytes. Representative assay from two independent experiments (3 biological replicates each) is shown; mean ± standard deviation. (D) Liver-stage activity was assessed using *P. berghei* as described;^9^ representative data from one of two independent studies (8 technical replicates each) shown; mean ± standard deviation The average EC_50_ across the two independent studies was EC_50_ = 0.57 ± 0.21 μM.

**Table 1:**
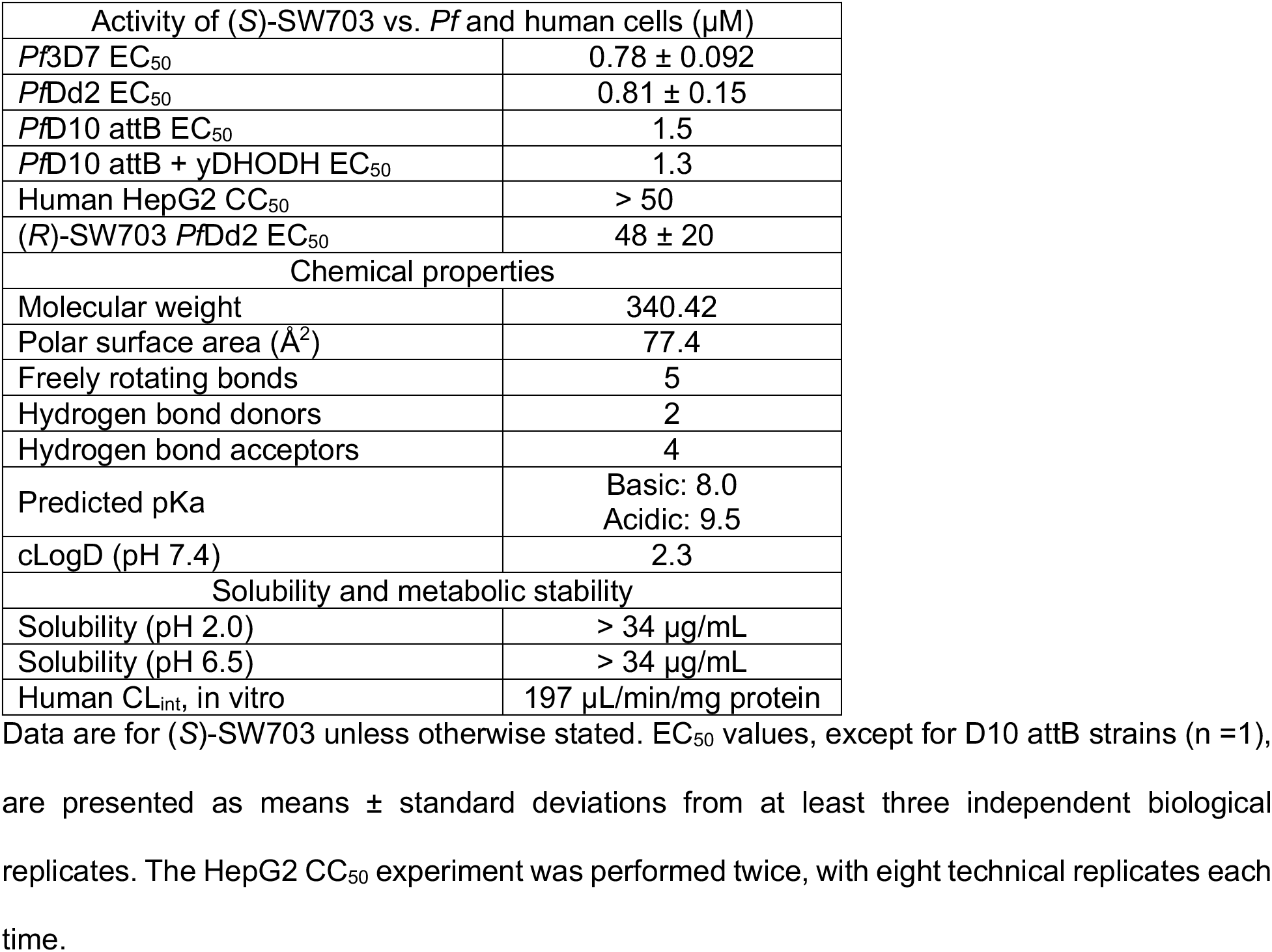
(*S*)-SW703 Cell activity, chemical properties and in vitro metabolism in human liver microsomes

(*S*)-SW703 contains a chiral center derived from L-Tyr and represents the S-enantiomer. Both (*S*)-SW703 and its R-enantiomer ((*R*)-SW703) were re-synthesized and tested for activity against cultured *P. falciparum* (asexual RBC stages) to determine whether the series shows enantiomeric selectivity. (*R*)-SW703 was found to be ~30-fold less active than the S-enantiomer (Dd2 EC_50_ = 48 ± 20 μM; Fig. 1b), suggesting that (*S*)-SW703 has a protein target.

### (*S*)-SW703 has fast-killing activity

Compounds with fast kill rates are of particular interest because they can provide rapid relief of malaria symptoms, which is especially important in cases of severe malaria; fast-killing compounds may also have a lower propensity for development of antimalarial resistance.^8, 26^ (*S*)-SW703 was tested to determine its killing kinetics by a previously described two-color FACS assay.^27^ Following 0, 24, or 48-hour exposure to each compound at 10X EC_50_, parasite viability was evaluated based on reinvasion of labeled RBCs (see Figure 1c). (*S*)-SW703 performed similarly to chloroquine and artesunate (benchmark antimalarials), consistent with a fast-killing mechanism.

### Activity of (*S*)-SW703 against liver-stage parasites and gamete formation

Liver-stage activity is an asset to potential novel antimalarial therapies, and is of particular value for chemoprophylaxis or prevention.^8^ (*S*)-SW703 liver-stage activity was assessed in a *P. berghei* liver stage model where infection is initiated in human HepG2 cells by incubation with sporozoites.^9^ HepG2 cells were pretreated with (*S*)-SW703 for 18 hours prior to infection with *P. berghei* sporozoites; parasite growth was measured 48 hours after infection. (S)-SW703 blocked propagation of the liver stage infection with similar potency to that observed against blood-stage parasites (EC_50_ = 0.57 ± 0.21 μM; see Figure 1d).

Transmission-blocking activity is also a desirable trait for novel antimalarials.^7, 8^ (*S*)-SW703 was tested for transmission-blocking activity against *P. falciparum* stage V gametocytes using the dual gamete formation assay (DGFA), in which cultured gametocytes (*P. falciparum* strain NF54) were incubated with 1 μM compound for 48 hours prior to induction of gamete formation.^28^ Male gamete formation was assessed immediately (exflagellation); female gamete formation was assessed after an additional 24 hours. At 1 μM (*S*)-SW703, 1.2% and 18.3% inhibition were observed for formation of male and female gametes, respectively. Given that the 1 μM concentration used in this assay is not substantially above the asexual blood stage EC_50_ (0.78 μM vs. *Pf*3D7; Table 1), the 18.3% inhibition of female gametes at 1 μM suggests that the series could have transmission-blocking activity, which may be sex-specific.

### Selection of (*S*)-SW703-resistant ((*S*)-SW703^R^) *PfDd2* parasites

In order to gain insight into the mechanism of action of (*S*)-SW703 against *P. falciparum,* we initiated selections to identify (*S*)-SW703 resistant parasites for whole genome sequencing (WGS). Two (*S*)-SW703 resistance screens were conducted: one used Dd2 as a parent strain; the other used genetically engineered Dd2 parasites with D308A and E301A mutations in the delta subunit of DNA polymerase (PfDd2_100022400; Dd2-Polδ); these changes increase the propensity for mutation.^29^ Both screens were performed using cycling drug selection method,^26, 30^ in which parasites were exposed to compound at ~10X EC_50_ for 48 h, or until all visible parasites were killed. Compound was then removed, and parasites were cultured in drug free media until parasites reemerged. Additional cycles were performed until resistant parasites were detected.

The Dd2-Polδ screen was performed using 3 flasks, each containing approximately 10^9^ parasites. After an initial 48-hour pulse at 5 μM (*S*)-SW703, parasites emerged in 9 days and the pulse was repeated once (2 pulses total), with parasites emerging 16 days after the end of the second pulse. Preliminary analysis on bulk cultures showed parasites had developed resistance to (*S*)-SW703 and limiting dilutions of recrudescent parasites were initiated in order to obtain clonal lines. Three clones from each screening flask were selected for WGS.

The Dd2 screen was performed using 4 flasks, each containing approximately 3 × 10^8^ parasites. Following an initial 48-hour pulse at 5 μM (*S*)-SW703, parasites recovered less than 7 days after compound was removed. The second pulse used 10 μM (*S*)-SW703 and was prolonged from 48 to 72 hours, because some live parasites were still observed by microscopy at 48 hours. After recovery, the third pulse (10 μM (*S*)-SW703) was initiated. The third pulse was elongated to 7 days (168 hours) because complete death of the culture was never observed. Compound was removed and clonal parasites were isolated by limiting dilutions; 3 clones from each screening flask were selected for resistance evaluation and WGS.

EC_50_ shifts in the (*S*)-SW703-resistant clones ranged from 2.2 to 58-fold relative to parental strains (Table 2). Selected EC_50_ curves for representative strains (one for each *Pf*CARL/*Pf*ACT mutation) are shown in Figure 2a. WGS analysis showed that all Dd2-Polδ resistant clones and 11 out of 12 Dd2 resistant clones have mutations in *PfCARL*(PfDd2_030027000; Table 2). The only (*S*)-SW703-resistant clone that did not contain a *Pf*CARL mutation had a Q116K point mutation in a putative acetyl coA transporter (*Pf*ACT; PfDd2_100041800). This mutation is in the last codon prior to a predicted intron and may therefore interfere with splicing; a splice defect would cause an insertion/frameshift, eventually ending in a premature stop codon and truncating the majority of the protein. *Pf*CARL and *Pf*ACT have previously been identified as multidrug resistance genes.^20, 21, 31–34^ Additionally, compounds with *Pf*CARL-mediated resistance mechanisms are often also subject to *Pf*ACT-mediated resistance.^21, 32^ Interestingly, (*S*)-SW703 EC_50_ shifts appeared to partition according to specific PfCARL mutations (Figure 2c): Q821H mutants are the most resistant (52 – 60 fold) relative to the parent strain, followed by L830I mutants (27 – 32 fold), S1057T mutants (12 – 16 fold), the single *Pf*CARL^L1073I^ mutant (9.5-fold resistance), and finally the *Pf*ACT^Q116K^ mutant (2.3-fold resistance). In representative parasite lines, the *Pf*CARL and *Pf*ACT mutations identified by the WGS analysis were verified by PCR of the *Pf*CARL or *Pf*ACT genes, followed by Sanger sequencing (Figures S1-S3, Table S1).

**Table 2:**
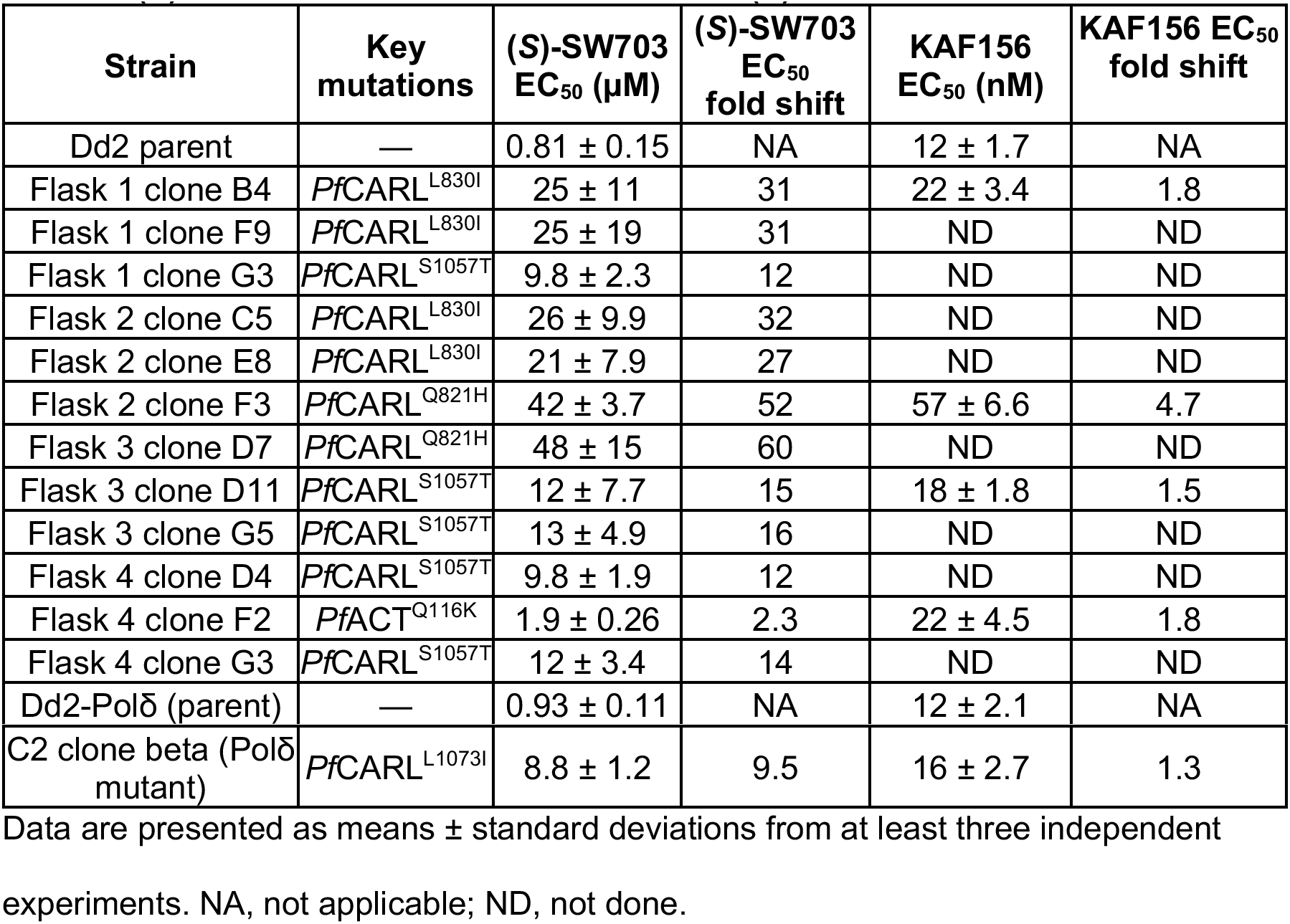
(*S*)-SW703 and KAF156 EC_50_ values for (*Ş*)-SW703^R^ clones.

**Figure 2.**
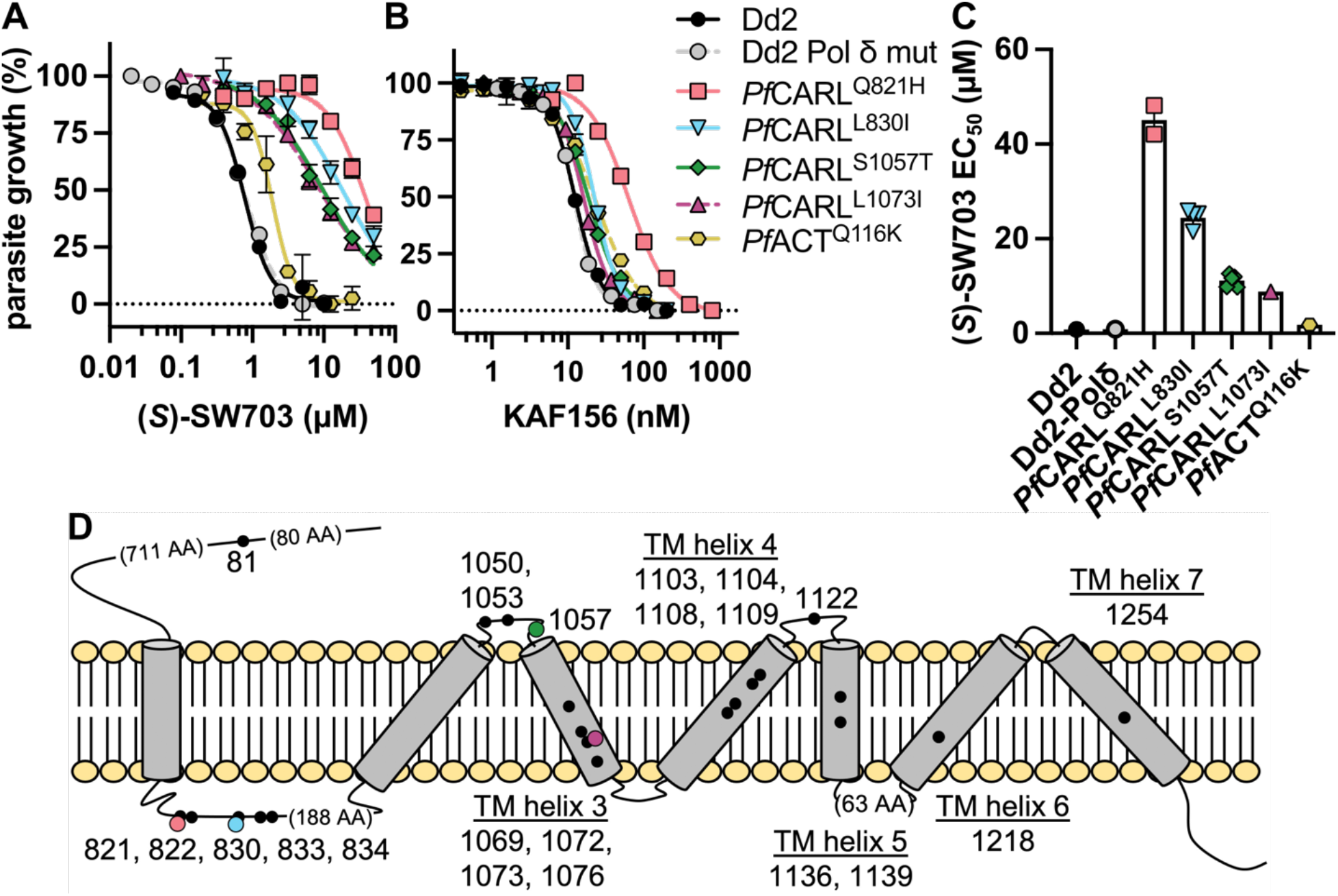
*Pf*CARL and *Pf*ACT mutations are associated with resistance to (*S*)-SW703. (A-B) Representative (*S*)-SW703 (A) and KAF156 (B) EC_50_ curves for parent strains and representative resistant clones; one clone for each *PfCARL* or *Pf*ACT mutation: *Pf*CARL^Q821H^, flask 2 clone F3; *Pf*CARL^L8301^, flask 1 clone B4; *Pf*CARL^S1057T^, flask 3 clone D11; *Pf*CARL^L1073I^, C2 clone beta; *Pf*ACT^Q116K^, flask 4 clone F2. Representative data from a single experiment are shown; mean ± standard deviation from three technical replicates. Averaged data from independent experiments (n ≥ 3) are presented in Table 2. (C) EC_50_ values of all clones for each *PfCARL* or *Pf*ACT mutation. Dd2 is the parent strain for *Pf*CARL^Q821H^, *Pf*CARL^L830I^, *Pf*CARL^S1057T^, and *Pf*ACT^Q116K^ clones; Dd2-Polδ mutant is the parent strain for *Pf*CARL^L1073I^. Datapoints represent mean EC_50_ values for each individual clone. Bars and error bars represent mean and standard deviation derived from these datapoints. (D) Cartoon model of *PfCARL* with locations of known mutations. Black dots represent previously reported mutations;^20, 21,31–35^ colored dots represent mutations identified in this study (Q821H, salmon pink; L830I, sky blue; S1057T, forest green; 1073I, plum purple). Note that in some cases, black dots represent multiple mutations at the same site. Mutations are labeled according to their amino acid positions; mutations that are predicted to lie within transmembrane helices (TMHMM^36^) are labeled above or below the helix.

Aside from changes to *PfCARL* and *Pf*ACT, other mutations of note identified by WGS included a mutation in multidrug resistance protein 1 (MDR1; *Pf*Dd2_050027900) at position 86. Mutations at MDR1 position 86 (in concert with others) have been associated with resistance to a variety of antimalarial compounds, especially those targeting the food vacuole (*Pf*MDR1 is localized mainly in the food vacuolar membrane).^37^ In 3D7, a drug-sensitive strain, the amino acid at position 86 is asparagine (N86). Dd2, the parent line used to initiate the selections, harbors several MDR1 mutations and amplifications, as compared to 3D7; in Dd2, the amino acid at position 86 is phenylalanine (F86). We observed a mutation to tyrosine (Y86). It is unclear what effect, if any, the F86Y mutation would have on (*S*)-SW703 resistance. The change was most prominent in one clone (flask 2 clone E8) from the Dd2 screen and all but one of the (*S*)-SW703^R^ clones from the Dd2-Polδ screen. However, Sanger sequencing consistently detected mixed populations of F86 and Y86 in MDR1, in all samples we sequenced (see Figure S3). All of these parasites had been cloned by limiting dilution, except that the Dd2-Polδ parental clone used for the screen was not available for sequencing (instead we sequenced an earlier sample from this strain). Our consistent observation of mixed populations, despite cloning for limiting dilutions, suggests that this position is highly mutable. Similar mixed alleles were seen for the *Pf*CARL^L1073I^ mutant, C2 clone beta, but the parental population appears clonal at this allele (Figure S1). We therefore suspect that either the limiting dilutions were insufficient to isolate single clones in this case, or that the mutation was unstable and the parasite population had already started to revert back to the wildtype. Copy number variation analysis was also performed but no notable copy number changes were detected (Table S3).

*Pf*CARL mutations were first identified in a screen for resistance to imidazolopiperazine compounds.^35^ Subsequent resistance screens using similar compounds have identified additional *Pf*CARL mutants,^20, 21, 31, 32^ and *Pf*CARL mutations have also been found in screens for resistance to additional unrelated scaffolds: MMV007564,^33^ and PPI28 compounds^34^ (Figure 3). At least 27 individual mutations in *Pf*CARL have been described to date (Figure 2D).^20, 21, 31–35^ Many of these mutations have been identified multiple times, in separate screens. While there is some overlap in resistance, some mutations appear to confer strong resistance to only specific compounds. Of the *Pf*CARL mutations identified in this study, *Pf*CARL^L830I^, *Pf*CARL^S1057T^, and *Pf*CARL^L1073I^ appear to be novel, although other mutations have previously been identified at positions 830 and 1073.^20, 31, 35^ The *Pf*CARL^Q821H^ mutation had previously been observed following selections for resistance to imidazolopiperazines or MMV007564.^20, 33, 35^

**Figure 3.**
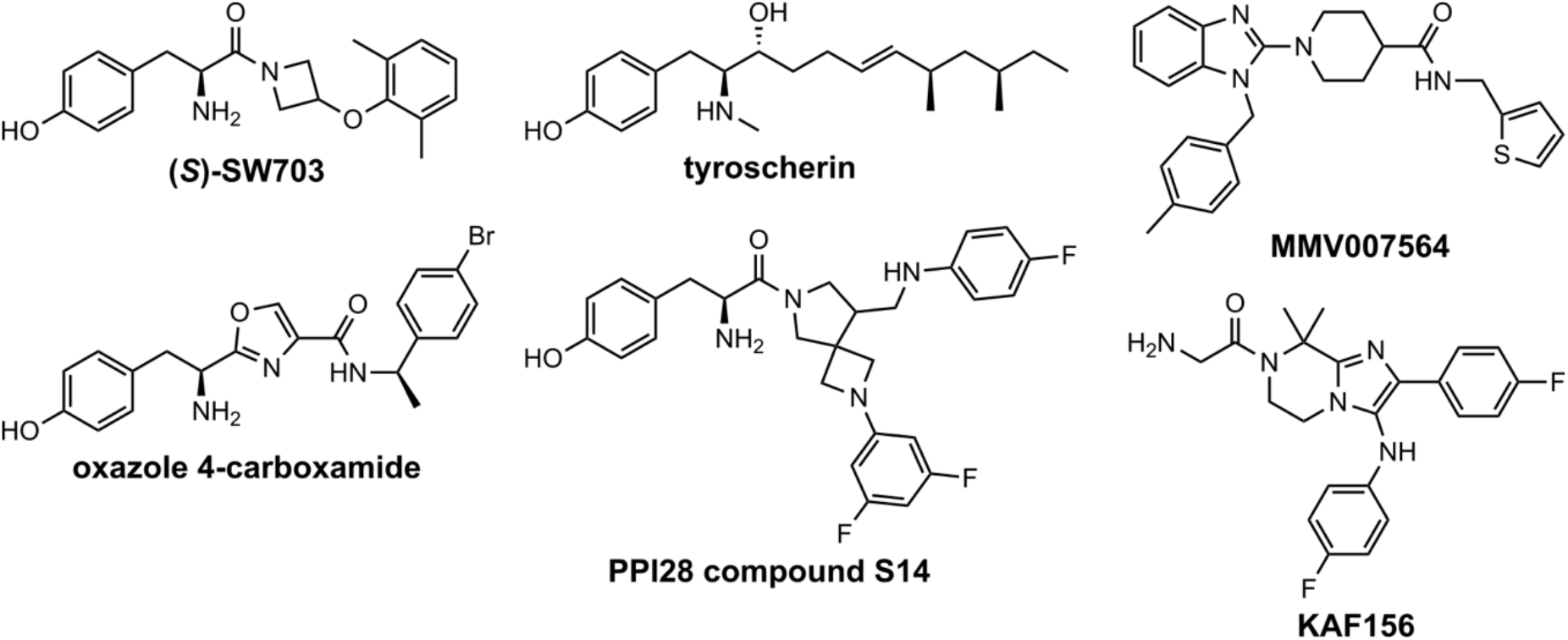
Structural comparison between (*S*)-SW703 and compounds reported to share the *Pf*CARL/*Pf*ACT resistance phenotype.^32, 34, 43^.

In light of the ongoing clinical development of KAF156, we tested our (*S*)-SW703-resistant clones to determine if they were cross-resistant to KAF156. Parasites bearing PfCARL^S1057T^, PfCARL^L830I^, PfCARL^L1073I^, and PfACT^Q116K^ mutations showed little (< 2X) cross-resistance to KAF156. Parasites bearing PfCARL^Q821H^ mutations showed about 5-fold resistance to GNF179 (a close analog of KAF156), similar to previous reports for this mutation^35^ (Figure 2B and Table 2). In contrast, very substantial (> 16X) cross resistance to (*S*)-SW703 was observed in a KAF156-selected PfCARL^I1139K^ mutant line ^35^ (Table 3). Other drug-resistant parasite strains generated to a range of preclinical and clinical candidates with different modes of action ^6, 17, 38, 39^ did not show any cross-resistance to (*S*)-SW703 (Table 3), suggesting that (*S*)-SW703 does not share a common target with these compounds. (*S*)-SW703 also showed equivalent activity against a set of clinical isolates that includes strains resistant to chloroquine (CQ), pyrimethamine (PYR), atovaquone (ATQ), and piperaquine (PPQ) ^40^ (Table 3).

**Table 3:**
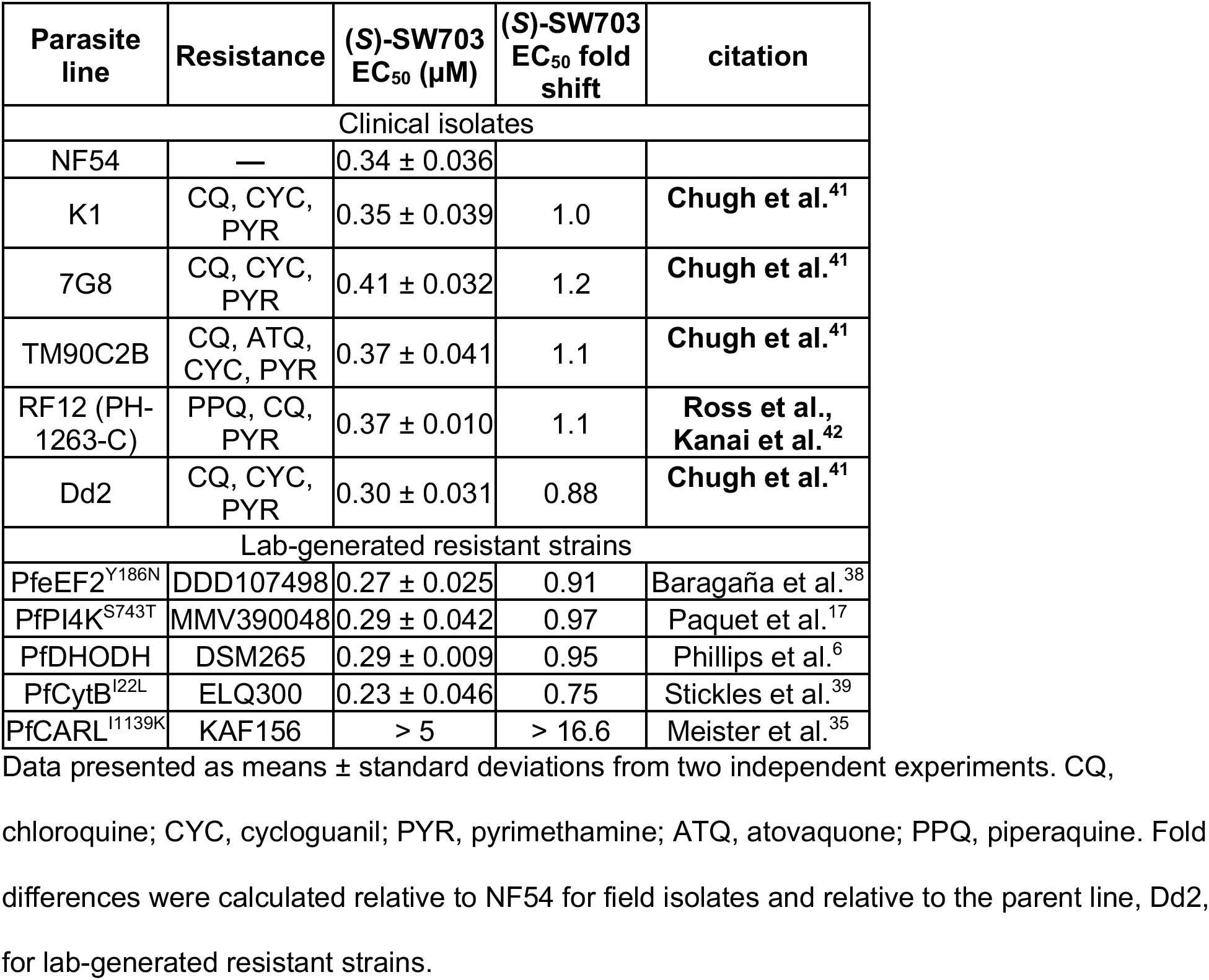
(*S*)-SW703 EC_50_ values for a panel of clinical isolates and lab-generated resistant strains

## Discussion

Malaria remains one of the most consequential global infectious diseases. The ability of the parasite to evade drug therapies through resistance threatens treatment and control programs and underscores the need for new antimalarial compounds to be identified and advanced to clinical development.^44^ Phenotypic HTSs have become an important tool in identifying new chemical classes for *P. falciparum* drug discovery.^11^ Using these methods, we identified (*S*)-SW703, a novel, fast-killing antimalarial with a tyrosine amide core and broad activity across different parasite life cycle stages. (*S*)-SW703 has low micromolar asexual blood-stage and liver-stage activity and our initial data also suggest that the series will have transmission-blocking activity versus female gametes, though confirmation of this activity would require more potent analogs. (*S*)-SW703 has good drug-like properties, excellent solubility, and was not cytotoxic to the human HepG2 cell line, providing an ideal starting point for further optimization. To obtain insight into the mechanism of (*S*)-SW703 killing, we used cycling drug pressure to select for resistant parasites. Whole genome sequencing revealed mutations in *Pf*CARL and *Pf*ACT, both which encode proteins associated with drug resistance to multiple unrelated chemical series. Although these studies did not directly elucidate a target and mechanism of action for (*S*)-SW703, they do identify a resistance mechanism primarily occurring through mutation of *Pf*CARL, providing a marker for resistance development.

Prior studies have reported that mutations in PfCARL were also associated with resistance to a number of unrelated scaffolds, including imidazolopiperidines such as KAF156—currently in phase II clinical development—and GNF179,^20, 21, 31, 32, 35^ MMV007564 (a benzimidazolyl piperidine),^33^ tyroscherin,^32^ an oxazole-4-carboxamide,^32^ and a diazaspiro[3.4]octanes series (PPI28).^34^ Mutations in *Pf*ACT and *Pf*UGT also confer multidrug resistance to many of these same scaffolds, suggesting that *Pf*CARL, *Pf*ACT and *Pf*UGT are multi-drug resistance proteins with a common resistance mechanism. ^21^

Although the precise mechanism of action has not been determined for any of the compounds that share the *Pf*CARL/*Pf*ACT/*Pf*UGT resistance mechanism, these proteins all localize to the ER (*Pf*ACT, *Pf*UGT) or cis-Golgi (*Pf*CARL),^21, 32^ and it has been suggested that the genes are resistance loci for compounds that act in the ER. Interestingly, PfCARL, PfACT, and PfUGT mutations slightly increased sensitivity to KDU691,^21, 32^ an inhibitor of *P. falciparum* phosphatidylinositol 4-kinase (PI4K), which is important for Golgi structure and function.^45^ Critical functions within the ER include lipid synthesis, calcium storage, synthesis of many proteins, post-translational modification, and initial transport of membrane proteins and trafficked proteins, many of which then transit through the Golgi. Most of these processes are thought to be essential throughout the parasite life cycle, consistent with the multi-stage potency of (*S*)-SW703 and other compounds. Functional studies of the imidazolopiperazines revealed that fluorescently-labeled GNF179 analogs accumulate in the ER, and that GNF179 causes defects in trafficked proteins, mild ER stress (eIF2α phosphorylation) and changes to ER morphology (expansion) in *P. falciparum*.^46^ ER-localized processes would therefore be of special interest in future efforts to identify the target(s) of (*S*)-SW703.

While compounds that share the *PfCARL* resistance mechanism are likely to act against ER-based targets, they may not act on the same targets within that compartment. Most of these chemotypes are distinct from each other, and from (*S*)-SW703 (see Figure 3). Rate of kill and activity against particular stages are both believed to correlate with mechanisms of action,^47^ and these compounds do not behave identically. (*S*)-SW703, the imidazolopiperazines, MMV007564, and the PPI28 compounds are all active against liver stage parasites. ^16, 20, 33, 34^ and most of the compounds have fast (PPI28 compounds, (*S*)-SW703) or fast/moderate (MMV007564) activity *in vitro*^27, 34^ or in vivo (imidazolopiperazines).^48^ However, reported differences between imidazolopiperazines and MMV007564 in stage specificity within the asexual intraerythrocytic development cycle have suggested that these compounds may have different targets.^33^ While (*S*)-SW703 shares its tyramine substructure with tyroscherin, the oxazole 4-carboxamide, and the PPI28 series (Fig. 3), (*S*)-SW703 has a strong enantiomeric selectivity that has not been observed in the PPI28 series. Most of the PPI28 compounds completely lack the amine moiety, and thus the stereocenter. For those PPI28 compounds that do retain this stereocenter, the enantiomers were shown to have roughly equivalent activity (e.g. S14 vs. 105),^34^ suggesting PPI28 and (*S*)-SW703 may have different targets and/or binding modes.

Of the PfCARL mutations that were found in (*S*)-SW703-resistant parasites, only Q821H has been observed previously. This mutation was identified as a resistance marker for both the imidazolopiperazines^35^ and MMV007564,^33^ and was associated with high level resistance in a double mutant with S1076R.^32 20^ However, while the leucine-to-isoleucine mutations we observed (L830I and L1073I) have not been reported, L830V was identified in at least four separate selections^20, 31–33, 35^ and confirmed by CRISPR mutagenesis to confer 20-to 35-fold resistance to KAF156.^32^ In contrast, our (*S*)-SW703-resistant PfCARL^L830I^ mutant was only 1.8-fold resistant to KAF156 (see Figure 2b and Table 2). L1073Q mutations were identified in selections with MMV007564^33^ and PPI28,^34^ though L1073Q did not confer resistance to KAF156 or GNF179.^33^ Similarly, our (*S*)-SW703-resistant PfCARL^L1073I^ clone was not resistant to KAF156. The last PfCARL mutation we observed in our (*S*)-SW703 selections, S1057T, appears to be completely novel. This observation suggests that (*S*)-SW703 may interact with PfCARL using an overlapping but unique binding mode compared to other compounds associated with PfCARL-based resistance mechanisms.

The only (*S*)-SW703-resistant clone that lacked mutations in PfCARL had a Q116K mutation in the acetyl-coA transporter (PfACT); PfACT had previously been identified as a multidrug resistance gene for many of the same compounds as PfCARL.^21^ Several point mutations (missense) and early stop mutations in PfACT showed high-level resistance to the imidazolopiperazine, GNF179 (> 92x EC_50_ shift) and this finding has been confirmed by generation of a CRISPR mutant (PfACT^S242^*), which conferred high-level resistance to both GNF179 and KAF156.^21^ In contrast, the Q116K mutant identified in our study had only very modest resistance towards not only (*S*)-SW703, but also KAF156. Since Q116 is the last codon before an intron, it is possible that this mutation disrupts splicing and results in an early stop codon, but the relatively modest impact of the mutation argues against that outcome.

## Conclusion

As the global malaria threat persists and grows, the need for novel treatments and prophylactic agents is dire. We discovered (*S*)-SW703, a tyrosine amide with rapid activity in asexual blood-stage parasites, equivalent potency against the liver-stage, and likely transmission-blocking activity. Resistance to (*S*)-SW703 is associated primarily with mutations in PfCARL and while the target of SW702 is unknown, this resistance profile suggests that (*S*)-SW703 is likely to act in the ER. A future candidate would ideally not be cross-resistant with any advanced molecule in the global portfolio. Given that *Pf*CARL and *Pf*ACT mutations were observed in (*S*)-SW703-resistant parasites, and the cross-resistance observed between (*S*)-SW703 and KAF156, this would need to be a goal of any future program on this chemical series. Nonetheless, the fast killing rate and the broad activity against life cycle stages, combined with the good drug-like properties and ease of synthesis, make the (*S*)-SW703 series an attractive starting point for the development of a novel antimalarial agent.

## Experimental

### Materials

The screening library of 100K compounds was obtained from Chemical Diversity Labs and ChemBridge Corporation. (*S*)-2-amino-1-(3-(2,6-dimethylbenzyl)azetidin-1-yl)-3-(4-hydroxyphenyl)propan-1-one ((*S*)-SW703) was purchased from Chembridge Corporation for hit validation work and subsequently synthesized for additional studies (see Supporting Information for synthetic methods and experimentals). DSM265 was on hand from prior work, synthesized as previously described.^6^ All compounds were at least 95% pure. Purity of (*S*)-SW703 from Chembridge Corporation and in-house synthesis was assessed by LC/MS; purity of DSM265 was assessed by HPLC.

### Methods

#### *P. falciparum* culture

Asexual blood-stage *P. falciparum* parasites were cultured at 37 °C, 5% CO_2_, using male O+ red blood cells (Valley Biomedical) in RPMI 1640 medium (Millipore Sigma), supplemented with 25 mM HEPES, 0.5% Albumax-I (Thermo Scientific), 23 mM sodium bicarbonate, 92 μM hypoxanthine, and 12.5 μg/mL gentamicin sulfate. Culture method was adapted from Trager and Jensen.^49^

### Small-molecule 100K library composition

A small-molecule library of 100K compounds belonging to the UT Southwestern HTS core was obtained from Chemical Diversity Labs and ChemBridge Corporation. The compounds in this collection obey Lipinski’s and Veber’s rules and do not contain PAINS substructures.^50^ The number of rings and stereo centers in each of the compounds have maximums of 4 and 3, respectively. Additionally, the compounds have QED^51^ (quantitative estimate for drug-likeness) >=0.5. All library compounds were dissolved in DMSO to a stock concentration of 5 mM.

### Compound library screening to identify *P. falciparum* growth inhibitors

Screening was conducted in the UT Southwestern HTS core. The *P. falciparum* growth inhibition assay was adapted from Smilkstein et al^22^ and followed the same screening protocol as previously published by our group for a smaller library.^23^ Prior to screening, 30 nL of compound (5 mM, DMSO) was added, per well, into 384-well plates using a Labcyte-Echo 555 dispenser. 60 μL of ring-stage *P. falciparum* parasites (pan-sensitive strain 3D7) were added to each well using a BioTex MultiFlo liquid dispenser, thus diluting the compounds to a final concentration of 2.5 μM compound (columns 3 – 22 of each plate). Pyrimethamine (0.5 μM) was used as a control for growth inhibition (column 1 of each plate); DMSO was used as a negative control (columns 2 and 23 of each plate). Parasites were incubated with compounds at 37 °C, 5% CO_2_. After 72 hours, plates were frozen at −80°C for at least 16 hours, then thawed at room temperature. 15 μL of concentrated lysis buffer (80 mM Tris pH 7.5, 20 mM EDTA, 0.32% saponin, 0.32% Triton X-100, 0.0012% SYBR green “10,000 x” solution (Sigma)) was added to each well and plates were incubated 4 hours in the dark at room temperature. Fluorescence was measured (485 nm excitation, 535 emission) using a BioTek Synergy H1 Hybrid plate reader. Z’ scores for each plate were calculated as Z’ = 1 – [(3 × DMSO standard deviation + 3 × pyrimethamine standard deviation)/absolute value (DMSO average – pyrimethamine standard deviation)]. RZ score refers to number of robust standard deviations (RSD) between the median measurement vs. the measurement for a particular compound, after correcting for plate, row, and column effects.^52^

### Hit Validation and counter screens in cytotoxicity assays

The top 300 hit compounds (RZ score < −7.6) were cherry picked and re-tested against both *Pf*3D7 (as described above) and a human leukemia cell line, K562 at 3 concentrations (3, 1, and 0.3 μM). Similar to parasite assays, compounds (30 nL) were dispensed into 384-well plates using a Labcyte-Echo 555 acoustic dispenser. K562 cells (60 μL, 600 cells per well) suspended in RPMI 1640 medium (Sigma) supplemented with 5% fetal bovine serum (heat inactivated, Gibco) and 2 mM glutamine (Gibco) were then added. CD437 (Sigma) and methotrexate were used as positive controls. Plates were incubated at 37 °C, 5% CO_2_. After 72 hours, viability was assessed using the CellTiter Glo assay (Promega), which measures intracellular ATP.

### Human HepG2 cells

To further evaluate cytotoxicity of (*S*)-SW703, cytotoxicity assays were also conducted versus HepG2 cells. Similar to parasite assays, compounds (30 nL) were dispensed into 384-well plates using a Labcyte-Echo 555 acoustic dispenser. HepG2 cells (60 μL, 600 cells per well) suspended in EMEM medium (Sigma) supplemented with 5% fetal bovine serum (heat inactivated, Gibco) and 2 mM glutamine (Gibco) were then added. CD437 (Sigma) and methotrexate were used as positive controls. Plates were incubated at 37 °C, 5% CO_2_. After 72 hours, viability was assessed using the CellTiter Glo assay (Promega), which measures intracellular ATP. Assays were performed twice; each biological replicate comprised technical triplicates.

### Two color flow cytometry rate of kill assay

The rate of kill assay was performed as in Linares et al.^27^ Experiments were carried out with strain 3D7A (BEI Resources, MRA-151, contributed by David Walliker). The assay was performed twice, with biological triplicates each time. The human biological samples were sourced ethically, and their research use was in accordance with the terms of the informed consents under an IRB/EC approved protocol

### Asexual blood-stage EC_50_ assays

After the initial screens, growth inhibition assays ((*S*)-SW703, (*R*)-SW703, and KAF156) were conducted in 384-well (60 μL per well) or 96-well (200 μL per well) plates. Compound stocks were prepared at 10 mM in DMSO; stock concentrations of KAF156 and DSM265 were further diluted to 50-200 μM and 100 μM, respectively. In all studies, DSM265 (0.8 – 100 nM) was plated as a control to ensure that its EC_50_ remained within a normal range (~5 – 10 nM). Compounds were added to the plate using a Tecan D300e automated low volume liquid handler. Ring-stage parasites (2% hematocrit, 0.5% parasitemia, final DMSO concentration 0.5%) were incubated with compounds, at a range of concentrations, for 72 hours at 37 °C, 5% CO_2_. Plates were frozen for at least 16 hours, then thawed at room temperature. For experiments performed in 384 well plates, 15 μL of concentrated SYBR Green lysis buffer (80 mM Tris pH 7.5, 20 mM EDTA, 0.32% saponin, 0.32% Triton X-100, 0.0012% SYBR Green “10,000 x” solution (Sigma)) was added and plates were incubated in the dark for 4 hours prior to reading. For experiments performed in 96 well plates, 100 μL of SYBR Green lysis buffer (20 mM Tris pH 7.5, 5 mM EDTA, 0.008% saponin, 0.2% Triton X-100, 0.0002% SYBR Green “10,000 x” solution (Sigma)) was added and well contents were mixed by pipetting. Fluorescence was measured (485 nm excitation, 535 or 528 nm emission) using a BioTek Synergy H1 Hybrid plate reader. Data were analyzed by nonlinear regression using GraphPad Prism (log(inhibitor) vs response—variable slope with four parameters). Experiments were performed with technical triplicates; data represent means ± standard deviations from at least 3 biological replicates. In cases where the EC_50_ exceeded 5-10 μM, growth inhibition was at the top of the range was insufficient to provide an accurate no-growth “plateau” for EC_50_ calculation. Therefore, the readout at 100 nM DSM265 (full inhibition) was used to define the bottom of the EC_50_ curve. Specifically, this technique was used to measure (*S*)-SW703 sensitivity for all of the (S)-SW703^R^ clones except for clone 4A (PfACT^Q116K^ mutant), and to estimate the EC_50_ of (*R*)-SW703. Cross-resistance testing with the panel of clinical isolates and lab-generated resistant strains was performed using the modified [^3^H]-hypoxanthine incorporation assay, as reported previously.^53^

### Liver stage activity assays

Liver stage activity assays, including follow-up assessment of toxicity to HepG2 host cells, were performed as in Swann et al.^9^ Assays were performed using 8 technical replicates. Data represent mean ± standard deviations from two biological replicates.

### Dual gamete formation assay (DGFA)

Dual gamete formation assay (DGFA, carryover format) was performed using 1 μM (*S*)-SW703, as prevoiusly.^13, 28^ Gametocytes were incubated with compound for 48 hours, after which male gamete formation was assessed by recording exflagellation using a Nikon TiE widefield microscope. After a further 24 hours, expression of Pfs25 on the surface of female gametes was assessed in a fluorescence assay using anti-Pfs25 conjugated to Cy3. Data are means from 4 biological replicates.

### ADME

Chemical properties (molecular weight, polar surface area, freely rotating bonds, hydrogen bond donors and acceptors, predicted pKa, and cLogD (pH 7.4)) were calculated using ChemAxon chemistry cartridge with JChem for Excel software (version 16.4.11).

For solubility measurements, compound was dissolved in DMSO and was spiked into phosphate buffer (pH 6.5) or 0.01 M HCl (approx. pH 2.0). The final DMSO concentration was 1%. After 30 minutes, solubility was determined by nephelometry analysis.^54^

For metabolic stability assays, 1 μM compound was incubated with human liver microsomes (Xenotech, lot# 1410230, 0.4 mg/mL protein) at 37 °C. An NADPH-regenerating system was added to initiate metabolism; control reactions did not contain NADPH.^55^ At various timepoints over the course of an hour, samples were collected and quenched using acetonitrile, with diazepam included as an internal standard. In vitro intrinsic clearance (CL_int_) was calculated from the apparent first-order degradation rate constant.

### Selection of (*S*)-SW703 resistant parasites

The Dd2-Polδ screen was performed using 3 flasks of parasites; each flask contained approximately 10^9^ parasites (80 mL per flask, 4% hematocrit, 3% parasitemia prior to treatment with compound, mostly ring-stage parasites). After an initial 48-hour pulse at 5 μM (*S*)-SW703, parasites were allowed to recover; cultures were monitored by microscopy. After recrudescence, the pulse was repeated once (2 pulses total). After the second recrudescence, clonal parasites were obtained from each flask by limiting dilutions.

The Dd2 screen was performed using 4 flasks, each containing approximately 3 × 10^8^ parasites (25 mL per flask, 4% hematocrit, 3% parasitemia at the beginning of each pulse, mostly ring-stage parasites). Pulse 1 (48 hours) was conducted at 5 μM (*S*)-SW703, pulse 2 required 10 μM (*S*)-SW703 over 72 hours to clear parasites, and pulse 3, also at 10 μM, failed to clear parasites, even after 7 days (168 hours), indicating that resistance had developed. Drug was removed and clonal parasites were obtained from each flask by limiting dilution. Clonal parasites from each flask were than analyzed for EC_50_ shifts as described above.

### Sample preparation for whole genome sequencing

For the Dd2 screen, genomic DNA samples were prepared from three clones from each of the four flasks used in screening, from the parental Dd2 clone, and from two “sister” clones of the Dd2 parent. For the Dd2-Polδ screen, genomic DNA samples were prepared from three clones from each of the three flasks used in screening, and from the parental Dd2-Polδ strain. For isolation of genomic DNA, infected RBCs were suspended in PBS and subjected to saponin treatment (0.03 – 0.1%) to lyse the RBCs. Genomic DNA was isolated from parasite pellets using Qiagen Blood and Cell Culture DNA Mini kit. Saponin-treated parasite pellets were first suspended in 200 μL PBS, as recommended for cultured cells. Two changes were made to the manufacturer’s instructions: 1) 400 ug (4 μL at 100 mg/mL) of RNase A was added along with the recommended proteinase K; 2) samples were eluted in water instead of the provided buffer.

### Whole genome sequencing (WGS) and analysis

Paired-end sequencing was performed with 150-base reads using Illumina NextSeq 500. WGS data are deposited in NCBI’s Sequence Read Archive (SRA) database (SRA BioProject ID PRJNA886692). The sequenced reads per sample were subjected to analysis for quality and contamination using FastQC v0.11.2^56^ and FastQ Screen v0.4.4,^57^ respectively. The reads from each sample were then mapped to the *Plasmodium falciparum* Dd2 reference genome using BWA-MEM,^58^ with default parameters. BWA efficiently aligns short reads sequences against a reference genome, allowing gaps and mismatches. Duplicated reads were marked and coverage was calculated using Picard tools (v1.127).

The potential SNPs and indels were discovered by running GATK’s (v3.5) HaplotypeCaller (parameters: -ploidy 1, --emitRefConfidence GVCF). Genotyping was then performed on samples pre-called with HaplotypeCaller using GenotypeGVCFs. Hard-filtering approach was used to filter variants. Briefly, SelectVariants module was used to subset SNPs and INDELS. Then VariantFilteration module was used to filter variants. Variants were annotated using SnpEff.^59^ Copy number variation analysis was performed using control-FREEC.^60^

### Sanger sequencing of PfCARL, PfACT, and PfMDR1 mutations

To verify mutations identified via WGS, regions of interest were PCR-amplified from genomic DNA and submitted for sequencing at the UT Southwestern Sanger Sequencing Core. Primers for amplification and sequencing are shown in Table S2.

## Supporting information

Table S3 - CNV results

## Supporting information

Experimental details of the synthetic methods, Tables S1 (full list of mutations observed in PfCARL, PfACT and PfMDR1) and S2 (cloning and sequencing primers), supplemental figures S1-S3 (Sanger sequencing traces for mutations in PfCARL, PfACT, and PfMDR1); PDF.

Table S3 (Copy number variation (CNV) information for wildtype parasites and (S)-SW703-resistant clones); XLSX.

## Author contributions

LSI performed the Dd2 resistance selection and subsequent EC_50_ assays, prepared samples for WGS and Sanger sequencing, and wrote the paper. AKL performed the initial HTS and confirmatory/cytotoxicity assays, performed the Dd2-Polδ resistance selection, and made key intellectual contributions to the project. SG synthesized (*S*)-SW703 and (*R*)-SW703. AK and CX performed mutational analyses of the WGS dataset. BL organized the profiling in MMV testing centers *(Plasmodium* lifecycle profiling, killing rate, resistant strains and clinical isolates) and evaluated the strength of the hits prioritized by the UT Southwestern team. NM, supervised by EAW, assessed liver stage activity. SW measured EC_50_ assays for the panel of resistant strains and clinical isolates. HN performed HTS. BP oversaw the HTS execution as well as data analysis and summarization. FJG and BCF assessed rate of killing (two-color flow cytometry assay). AC, supervised by JB, performed the DGFA assay (transmission-blocking activity). JMR evaluated the screening hits for good chemical properties and supervised synthetic chemistry. MAP conceived and supervised the parasitology and overall project and wrote the paper.

## Acknowledgement

We are grateful to Dr. Marcus Lee and Dr. Krittikorn Kumpornsin (Wellcome Sanger Institute) for the gift of the Dd2-Polδ strain used in one of our (*S*)-SW703 resistance screens. We thank Dr. Anwu Zhou for processing the data for the primary screen and hit confirmation studies. The authors would like to acknowledge the use of the PlasmoDB, *Plasmodium* informatics resource, as part of the work of this paper. Dr. Susan Charman at the Centre for Drug Candidate Optimisation (CDCO), Monash University is acknowledged for conducting the ADME studies. We thank Christian Scheurer for technical assistance with the *P. falciparum* in vitro efficacy studies performed with the panel of clinical isolates and lab-generated resistant strains.

This work was funded in part by funds from the United States National Institutes of Health grants, R01AI103947 (to MAP), R01AI155784 (to MAP and JMR), and NIGMS predoctoral training grant GM007062 (to AL), and by Medicines for Malaria Venture (MMV) through their assay support network to JB (RD-08-2800), EW, SW, and FJG. JB acknowledges support from Wellcome via in Investigator Award (100993/Z/13/Z). BAP acknowledges the support of the HTS Core facility by U. T. Southwestern Medical Center and the State of Texas. The CDCO is funded in part by the Monash University Technology Research Platform network and Therapeutic Innovation Australia (TIA) through the Australian Government National Collaborative Research Infrastructure Strategy (NCRIS) program. MAP and JMR also acknowledge the support of the Welch Foundation (I-1257, I-1612). MAP holds the Sam G. Winstead and F. Andrew Bell Distinguished Chair in Biochemistry.

## Abbreviations used

ATQ: atovaquone
BRRoK: bioluminescence relative rate of kill
CLint: intrinsic clearance
CQ: chloroquine
CYC: cycloguanil
Dd2-Polδ: Dd2 with mutant DNA polymerase δ
DGFA: dual gamete formation assay
DMSO: dimethylsulfoxide
eIF2α: eukaryotic translation initiation factor 2 alpha subunit
ER: endoplasmic reticulum
FACS: fluorescence-activated single cell sorting
IGF: insulin-like growth factor
MDR1: multidrug resistance protein 1
MMV048: MMV390048
PfACT: *P. falciparum* acetyl-coA transporter
PfCARL: *P. falciparum* cyclic amine resistance locus
PfUGT: *P. falciparum* UDP-galactose transporter
PI4K: phosphatidylinositol 4-kinase
PPQ: piperaquine
PYR: pyrimethamine
RBCs: red blood cells
SD: standard deviation
(*S*)-SW703: (*S*)-SW228703
WT: wildtype.

## For Table of Contents Only

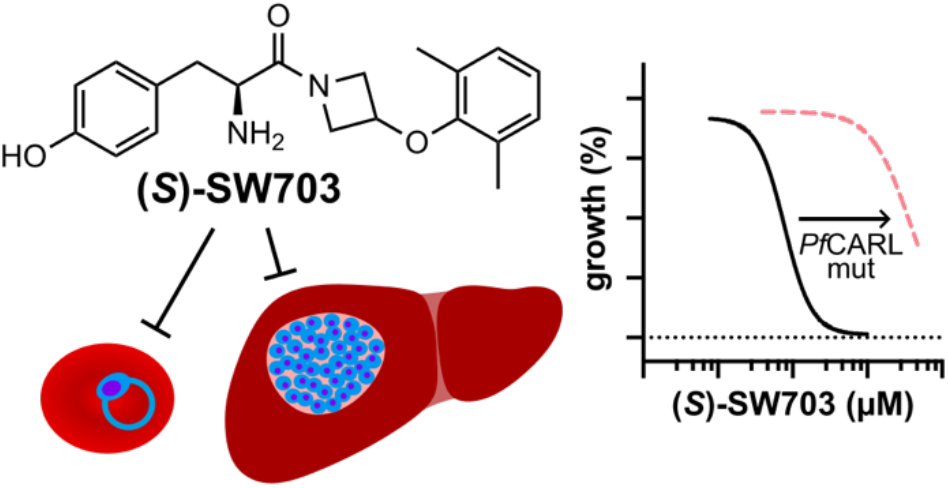

