## Supplemental methods, tables and figures for "A fast-killing tyrosine amide ((*S*)-SW228703) with blood and liver-stage antimalarial activity associated with the Cyclic Amine Resistance Locus (*Pf*CARL)"

[1]Department of Biochemistry; [2]Eugene McDermott Center for Human Growth and Development; [3]Department of Bioinformatics, University of Texas Southwestern Medical Center, Dallas, Texas, United States; [4]Division of Host-Microbe Systems and Therapeutics, Department of Pediatrics, UC San Diego; [5]Swiss Tropical and Public Health Institute; [6]University of Basel, Switzerland; [7]Department of Life Sciences, Imperial College London; [8]Global Health Medicines, GSK R&D, Tres Cantos, Spain; [9]Medicines for Malaria Venture, 1215 Geneva, Switzerland; [10]School of Biomedical Sciences, University of New South Wales, Sydney, Australia

*corresponding author

**Supplemental Experimental Methods.**

***Chemical synthesis of (S)-SW703 and (R)-SW703.***

**General Considerations**. All reactions were performed with commercially obtained anhydrous solvents. All reactions and final products were monitored by liquid chromatography mass spectrometry (LC/MS) with an Agilent Technologies 1200 series LC/MS using electrospray ionization methods. $^{1}H$ and $^{13}C$ NMR spectra were recorded on Bruker 400 MHz, or 600 MHz spectrometer. Chemical shifts $\delta$ are in ppm and spectra were referenced using the residual solvent peak. The following abbreviations are used: singlet (s), doublet (d), triplet (t), quartet (q), double doublet (dd), quintet (quin), multiplet (m), broad signal (bs). Final compounds were assessed to be at least 95% pure.

***tert*-butyl (*S*)-(1-(3-(2,6-dimethylphenoxy)azetidin-1-yl)-3-(4-hydroxyphenyl)-1-oxopropan-2-yl)-$\lambda^2$-azanecarboxylate (3)**

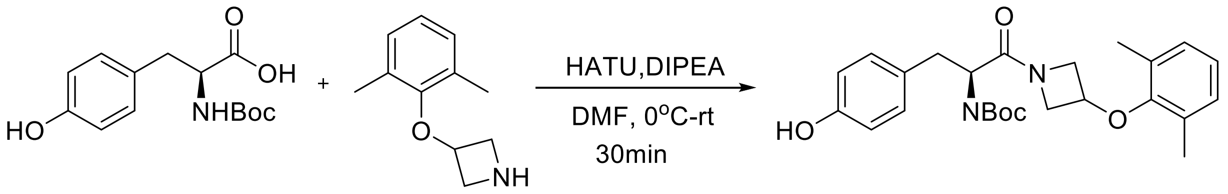


**1** **2** **3**

To a solution of the *L*-Boc-Tyr **1** (79 mg, 0.28 mmol), 3-(2,6-dimethylphenoxy)azetidine (**2**) (50 mg, 0.28 mmol) and HATU (129 mg, 0.34 mmol) in dry DMF (2 mL) was added DIPEA (0.19 mL, 1.12 mmol) at 0 ºC. The reaction mixture was then stirred for 30 min at room temperature. After that the mixture was diluted with ethyl acetate, washed with water and brine. The organic layer was dried over anhydrous $Na_2SO_4$, and the solvent was evaporated under vacuum. The residue obtained was purified by flash column chromatography (EtOAc/ hexane 3 : 2 v/v) to furnish the coupled product *tert*-butyl (*S*)-(1-(3-(2,6-dimethylphenoxy)azetidin-1-yl)-3-(4-hydroxyphenyl)-1-oxopropan-2-yl)-l2-azanecarboxylate (**3**) (110 mg, 90%) as colorless liquid.

$^1$H NMR (400 MHz, MeOD) $\delta$ 7.12 - 7.01 (m, 2H), 6.99 - 6.93 (m, 2H), 6.91 - 6.84 (m, 1H), 6.81 - 6.75 (m, 1H), 6.75 - 6.67 (m, 1H), 4.56 - 4.48 (m, 0.5H), 4.42 - 4.01 (m, 4H), 3.89 (dd, *J* = 10.9, 4.8 Hz, 0.5H), 3.75-3.69 (m, 0.5H), 3.44 (dd, *J* = 9.6, 4.7 Hz, 0.5H), 2.95 - 2.72 (m, 2H), 2.10 (s, 3H), 2.09 (s, 3H), 1.42 (d, *J* = 6.3 Hz, 9H).

$^{13}$C NMR (151 MHz, MeOD) $\delta$ 173.7 (d), 157.7 (d), 157.4, 156.6 (d), 131.4 (d), 131.3, 129.9 (d), 128.4 (d), 125.3 (d), 116.3 (d), 80.6, 71.4 (d), 59.6 (d), 57.1 (d), 53.7 (d), 38.4 (d), 28.6, 17.0 (d).

ESI-MS (m/z): 441.2 [M+H]+.

**(*S*)-2-amino-1-(3-(2,6-dimethylphenoxy)azetidin-1-yl)-3-(4-hydroxyphenyl)propan-1-one ((*S*)-SW703)**

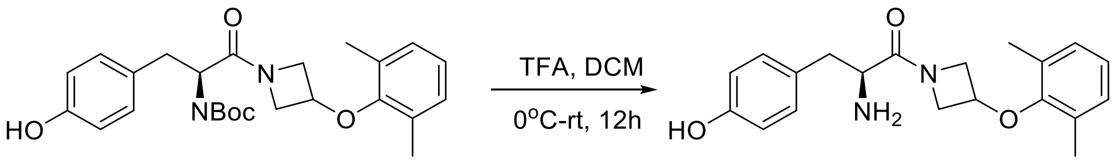


**3**

To a solution of the Boc-protected amine from the previous step (**3**) (40 mg, 0.09 mmol) in dry DCM (2 mL) was added TFA ( 0.14 mL, 1.81 mmol) at 0 ºC, and the resulting reaction mixture was stirred for 6 h at room temperature. The solvent was evaporated under vacuum. Then the reaction mixture was neutralized with saturated aqueous $NaHCO_3$ and extracted with DCM. The combined organic layers were washed with brine, dried over anhydrous $Na_2SO_4$ and concentrated under vacuum. The crude product was purified by flash column chromatography using ($CH_3OH/CH_2Cl_2$ 1 : 19 v/v) as eluent to give title compound **(*S*)-SW703** (24 mg, 81%) as a white solid.

$^{1}H$ NMR (400 MHz, $CD_3OD$) $\delta$ 7.10 - 6.99 (m, 2H), 6.99 - 6.93 (m, 2H), 6.94 - 6.84 (m, 1H), 6.82 - 6.75 (m, 1H), 6.75 - 6.68 (m, 1H), 4.58 - 4.45 (m, 0.5H), 4.27 - 4.11 (m, 2.5H), 4.08 - 4.02 (m, 0.5H), 3.93 - 3.87 (m, 0.5H), 3.55 - 3.41 (m, 1.5H), 3.30-3.26 (m, 0.5H), 2.86 - 2.77 (m, 1H), 2.75 - 2.65 (m, 1H), 2.13 (s, 3H), 2.10 (s, 3H).

$^{13}C$ NMR (101 MHz, MeOD) $\delta$ 175.9 (d), 157.7 (d), 156.7 (d), 131.3 (d), 131.2, 129.9 (d), 129.1(d), 125.3 (d), 116.3 (d), 71.5 (d), 59.4 (d), 57.0 (d) , 54.1(d), 42.4(d), 17.0(d).

ESI-MS (m/z): 341.2 [M+H]+.

### Supplemental Tables

#### Table S1: Full list of mutations observed in (*S*)-SW703-resistant parasites in *Pf*CARL, *Pf*ACT and *Pf*MDR1

| Strain | ***Pf*CARL (PfDd2_030027000)** | ***Pf*ACT (PfDd2_100041800)** | ***Pf*MDR1[a] (PfDd2_050027900)** |
|---|---|---|---|
| Dd2 screen | | | |
| Dd2 parent (clone G9) | Q821, L830, S1057 | Q116 | F86 |
| Flask 1 clone B4 | **L830I** | Q116 | F86 |
| Flask 1 clone F9 | **L830I** | Q116 | F86 |
| Flask 1 clone G3 | **S1057T** | Q116 | F86 |
| Flask 2 clone C5 | **L830I** | Q116 | F86 |
| Flask 2 clone E8 | **L830I** | Q116 | **F86Y** |
| Flask 2 clone F3 | **Q821H** | Q116 | F86 |
| Flask 3 clone D7 | **Q821H** | Q116 | F86 |
| Flask 3 clone D11 | **S1057T** | Q116 | F86 |
| Flask 3 clone G5 | **S1057T** | Q116 | F86 |
| Flask 4 clone D4 | **S1057T** | Q116 | F86 |
| Flask 4 clone F2 | Q821, L830, S1057 | **Q116K** | F86 |
| Flask 4 clone G3 | **S1057T** | Q116 | F86 |
| Dd2-Polδ screen | | | |
| Dd2-Polδ (parent)[b] | Q821, L830, S1057 | Q116 | F86 |
| C1 clone alpha | **L1073I** | Q116 | **F86Y** |
| C1 clone beta | **L1073I** | Q116 | **F86Y** |
| C1 clone gamma | **L1073I** | Q116 | **F86Y** |
| C2 clone alpha | **L1073I** | Q116 | **F86Y** |
| C2 clone beta | **L1073I**[c] | Q116 | **F86Y** |
| C2 clone gamma | **L1073I** | Q116 | **F86Y** |
| C3 clone alpha | **L1073I** | Q116 | F86 |
| C3 clone beta | **L1073I** | Q116 | **F86Y** |
| C3 clone gamma | **L1073I** | Q116 | **F86Y** |

Mutations (SNPs, bold) were originally detected by WGS and these changes were confirmed by Sanger sequencing in all parental and clonal lines from the Dd2 screen, and for the C2 clone beta clone from the Dd2-Polδ screen; see Figures S1-S3. WGS data are available under SRA BioProject ID PRJNA886692.

[a]Sanger sequencing of *Pf*MDR1 (PFDd2_050027900) showed mixed F86 and Y86 alleles for all parasites sequenced (see Figure S1). This table shows results as called during WGS analysis.

[b]The Dd2-Polδ screen was performed using a clone from recent limiting dilutions; however, the WGS and Sanger sequencing of the parental line were performed using gDNA from a prior population that was possibly more mixed.

[c]Sanger sequencing of *Pf*CARL (PFDd2_030027000) showed mixed L1073 and I1073 alleles for C2 clone beta (see Figure S1).

**Table S2. Primers for amplification and Sanger sequencing**

| Gene | Positions of interest | Primers |
|---|---|---|
| *Pf*CARL (PfDd2_030027000) | 821 (Q821H), 830 (L830I) | F: 5’-GAACGAAAATAATAAGAAGGATTCA-3’<br>R: 5’-ACTAGATTTTTCGTTATAGTTATTTCT-3’ |
| *Pf*CARL (PfDd2_030027000) | 1057 (S1057T), 1073 (L1073I) | F: 5’- ATCATGCAGATTCATTTTATTATCAG-3’<br>R: 5’- AACAGAACAAGAAAAGCATCTATG-3’ |
| *Pf*ACT (PfDd2_100041800) | 116 (Q116K) | F: 5’-CCAATTGGAGTTTCATCCGTAG-3'<br>R: 5’-GTTCGGAATATTGTCTTTTTCTCC-3' |
| *Pf*MDR1 (PfDd2_050027900) | 86 (F86Y) | F: 5’-GAGTACCGCTGAATTATTTAGAA-3'<br>R: 5’-TTATTATCATGAAATTGTCCATCTTG-3' |

### Supplemental Figures

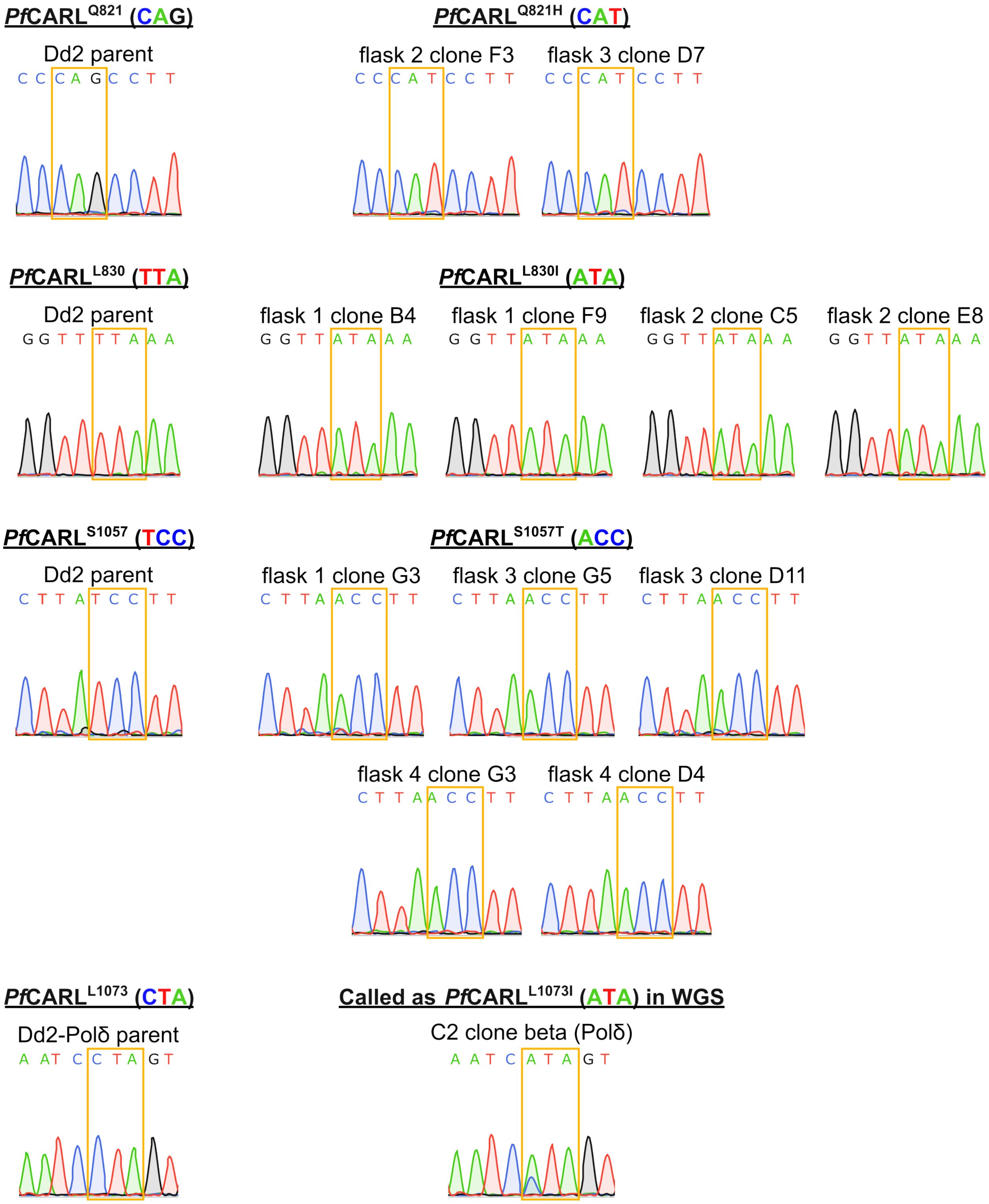


**Figure S1.** Sanger sequencing to verify mutations in *Pf*CARL (PfDd2_030027000).

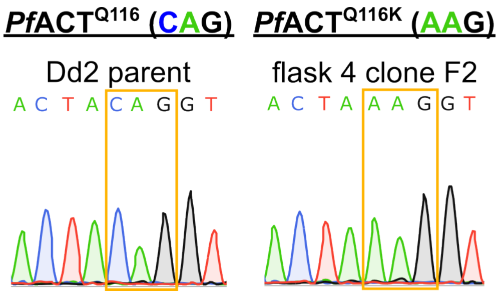


**Figure S2.** Sanger sequencing of *Pf*ACT (PfDd2_100041800) to verify the Q116K mutation.

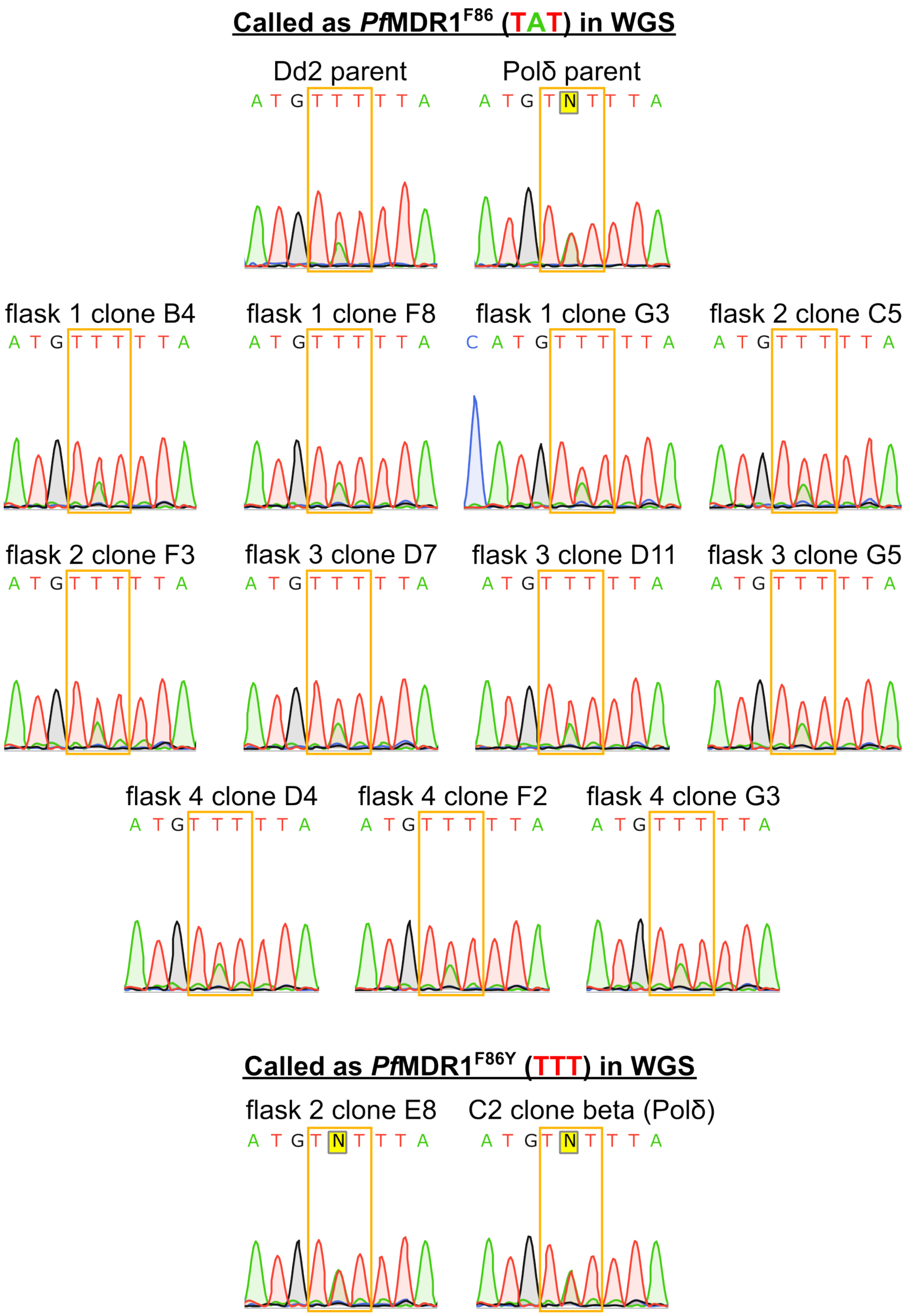


**Figure S3.** Sanger sequencing of *Pf*MDR1 (PfDd2_050027900) showed a mixture of F86 and Y86 alleles for all samples.
